# LL-37 fights SARS-CoV-2: The Vitamin D-Inducible Peptide LL-37 Inhibits Binding of SARS-CoV-2 Spike Protein to its Cellular Receptor Angiotensin Converting Enzyme 2 *In Vitro*

**DOI:** 10.1101/2020.12.02.408153

**Authors:** Annika Roth, Steffen Lütke, Denise Meinberger, Gabriele Hermes, Gerhard Sengle, Manuel Koch, Thomas Streichert, Andreas R. Klatt

## Abstract

**Objective:** Severe acute respiratory syndrome coronavirus-2 (SARS-CoV-2) is the pathogen accountable for the coronavirus disease 2019 (COVID-19) pandemic. Viral entry via binding of the receptor binding domain (RBD) located within the S1 subunit of the SARS-CoV-2 Spike (S) protein to its target receptor angiotensin converting enzyme (ACE) 2 is a key step in cell infection. The efficient transition of the virus is linked to a unique protein called open reading frame (ORF) 8. As SARS-CoV-2 infections can develop into life-threatening lower respiratory syndromes, effective therapy options are urgently needed. Several publications propose vitamin D treatment, although its mode of action against COVID-19 is not fully elucidated. It is speculated that vitamin D’s beneficial effects are mediated by up-regulating LL-37, a well-known antimicrobial peptide with antiviral effects.

**Methods:** Recombinantly expressed SARS-CoV-2 S protein, the extended S1 subunit (S1e), the S2 subunit (S2), the receptor binding domain (RBD), and ORF8 were used for surface plasmon resonance (SPR) studies to investigate LL-37’s ability to bind to SARS-CoV-2 proteins and to localize its binding site within the S protein. Binding competition studies were conducted to confirm an inhibitory action of LL-37 on the attachment of SARS-CoV-2 S protein to its entry receptor ACE2.

**Results:** We could show that LL-37 binds to SARS-CoV-2 S protein (LL-37/S_Strep_ K_D_ = 410 nM, LL-37/S_His_ K_D_ = 410 nM) with the same affinity, as SARS-CoV-2 binds to hACE2 (hACE2/S_Strep_ K_D_ = 370 nM, hACE2/S_His_ K_D_ = 370 nM). The binding is not restricted to the RBD of the S protein, but rather distributed along the entire length of the protein. Interaction between LL-37 and ORF8 was detected with a K_D_ of 290 nM. Further, inhibition of the binding of S_Strep_ (IC_50_ = 740 nM), S1e (IC_50_ = 170 nM), and RBD (IC_50_ = 130 nM) to hACE2 by LL-37 was demonstrated.

**Conclusions:** We have revealed a biochemical link between vitamin D, LL-37, and COVID-19 severity. SPR analysis demonstrated that LL-37 binds to SARS-CoV-2 S protein and inhibits binding to its receptor hACE2, and most likely viral entry into the cell. This study supports the prophylactic use of vitamin D to induce LL-37 that protects from SARS-CoV-2 infection, and the therapeutic administration of vitamin D for the treatment of COVID-19 patients. Further, our results provide evidence that the direct use of LL-37 by inhalation and systemic application may reduce the severity of COVID-19.

## Introduction

Severe acute respiratory syndrome coronavirus type 2 (SARS-CoV-2) is one of seven known human coronavirus pathogens ^1^. Most human coronavirus infections result in the common cold and account for up to 15% of such cases. In contrast, SARS-CoV, MERS-CoV, and SARS-CoV-2 infections can develop into life-threatening lower respiratory syndromes ^2–4^. As of November 28, 2020, a total of 61,508,373 infected cases and more than 1,441,265 deaths (mortality ~2%) were reported (WorldOmeter https://www.worldometers.info/coronavirus/). Coronavirus disease 2019 (COVID-19) has resulted in a global pandemic. An end of the pandemic will be reached if a majority of the population becomes immune after surviving the infection or through an effective vaccine. In addition to vaccine development, research focuses on therapy options. Attractive therapeutic approaches are the inhibition of SARS-CoV-2 fusion and entry, the disruption of virus replication, the suppression of excessive inflammatory response, and the treatment with convalescent plasma ^5^.

The COVID-19 mortality rate is currently higher at northern latitudes, a possible role for vitamin D (1,25(OH)_2_D) in determining outcomes has been postulated ^6^. Vitamin D deficiency is considered a global problem and might be associated with increased susceptibility to COVID-19. A recent study demonstrated that administration of a high dose of 25-hydroxyvitamin D (25(OH)D), the main metabolite of vitamin D, significantly reduces the need for intensive care unit (ICU) treatment of patients requiring hospitalization due to proven COVID-19 ^7^. One of 50 patients treated with 25(OH)D required admission to the ICU (2%), in comparison to 13 of 26 untreated patients required admission to the ICU (50%). Vitamin D might therefore be a promising therapeutic agent for fighting SARS-CoV-2. High‐dose vitamin D supplementation should achieve and maintain optimal (range 40‐60 ng/ml) serum 25(OH)D levels both for COVID-19 prevention and treatment ^8^.

Vitamin D is not only known for its role in bone mineralization and calcium homeostasis, but is also involved in various pathophysiological mechanisms that occur during SARS-CoV-2 infection. It was suggested, that vitamin D increases the ratio of angiotensin converting enzyme (ACE)2 to ACE, thus increasing angiotensin II hydrolysis and reducing subsequent inflammatory cytokine response to pathogens and lung injury ^6^. Also, many studies show that vitamin D has a decisive influence on the immune system and regulates antimicrobial innate immune responses ^9–11^. The effects of vitamin D are mediated by its binding to the vitamin D receptor (VDR) and subsequent binding to vitamin D response elements (VREs) ^9^. Such a consensus VRE is located in the promotor of the human cathelicidin antimicrobial peptide gene. Therefore, cathelicidin, the precursor of the well investigated human antimicrobial peptide LL-37, is strongly up-regulated by vitamin D ^12,13^.

LL-37 consists of 37 amino acids with an overall positive net charge (+6) and can thus eliminate microbes directly by electrostatic binding to negatively charged molecules on microbial membranes. In addition, it has antiviral effects including inhibition of herpes simplex virus type one (HSV-1) replication, vaccinia virus replication, retroviral replication, and replication of some adenovirus serotypes ^14^. The epidemiological evidence for a positive vitamin D-related immune effect includes many studies that feature enveloped viruses. This supports the notion that LL-37’s antiviral effects may be partially mediated by envelope disruption ^15^. SARS-CoV-2 is an enveloped single-stranded RNA virus harboring a genome of approximately 30 kb that encodes, among others, for the structural spike (S) glycoprotein that forms trimers at the viral surface. The S protein is important for viral attachment to the host cell via ACE2 and comprises two subunits, the S1 subunit containing the receptor binding domain (RBD) and the S2 subunit ^16^. Furthermore, the genome codes for a total of six accessory proteins including open reading frame (ORF)8, which might function as a potential immune modulator to delay or attenuate the host immune response against the virus ^17^. Interestingly, the S protein and the accessory protein ORF8 have a negative net charge of −8 and −4 (https://pepcalc.com/), respectively. Therefore, we speculated that LL-37 interacts with SARS-CoV-2 S protein and ORF8 and block viral entry.

In this study, we investigated the interaction between LL-37 and the SARS-CoV-2 S protein and ORF8 protein by surface plasmon resonance (SPR). Furthermore, we examined whether LL-37 inhibits the binding of the S protein to its cellular receptor hACE2.

## Material and Methods

### Cloning and Purification of SARS-CoV-2 proteins

The genes encoding ectodomain from the spike glycoprotein (MN908947; AA:1-1207; furin site mutated; K986P and V987P) and the S2 subunit (MN908947; AA:686-1207; K986P and V987P) including a C-terminal T4 foldon, the extended S1 subunit (MN908947; AA:14-722; furin site mutated) and the receptor binding domain (MN908947; AA:331-524) regions of the SARS-CoV-2 spike DNA as well as the ORF8 (MN908947; AA:16-121) and the human ACE2 (NM_001371415.1: AA:20-611) were amplified from synthetic gene plasmids using specific PCR-primers. After digestion with appropriate restriction enzymes the PCR products were cloned into modified sleeping beauty transposon expression vectors ^18^ that contain a double Strep II or a poly-histidine tag (Figure 1 A) and a thrombin cleavage site at the C-terminus. In the case of RBD, S1e, ORF8, and ACE2 the tag was added at the 5’ end including a BM40 signal peptide sequence. The sleeping beauty transposon system was stably transfected into HEK293 EBNA cells using FuGENE^®^ HD (Promega GmbH, USA) in DMEM/F12 (Merck, Germany) supplemented with 6% FBS (Biochrom AG, Germany). Cells were quickly selected with puromycin dihydrochloride (3 μg/ml; Sigma, USA) for four days and seeded into triple flasks. After four days of induction with doxycycline hyclate (0.5 μg/ml; Sigma, USA) supernatants were harvested every third day and filtered. Recombinant proteins with a double Strep II tag were purified using Strep-Tactin^®^XT (IBA Lifescience, Germany), eluted with biotin elution buffer (IBA Lifescience, Germany). In the case of poly-histidine-tagged proteins, cell supernatants were purified via an Indigo-Ni column (Cube Biotech, Germany), washed step-wise with imidazole, and eluted with 200 mM imidazole. Proteins were dialyzed against Tris-buffered saline (TBS) or phosphate-buffered saline (PBS) and aliquots stored at 4°C.

**Figure 1:**
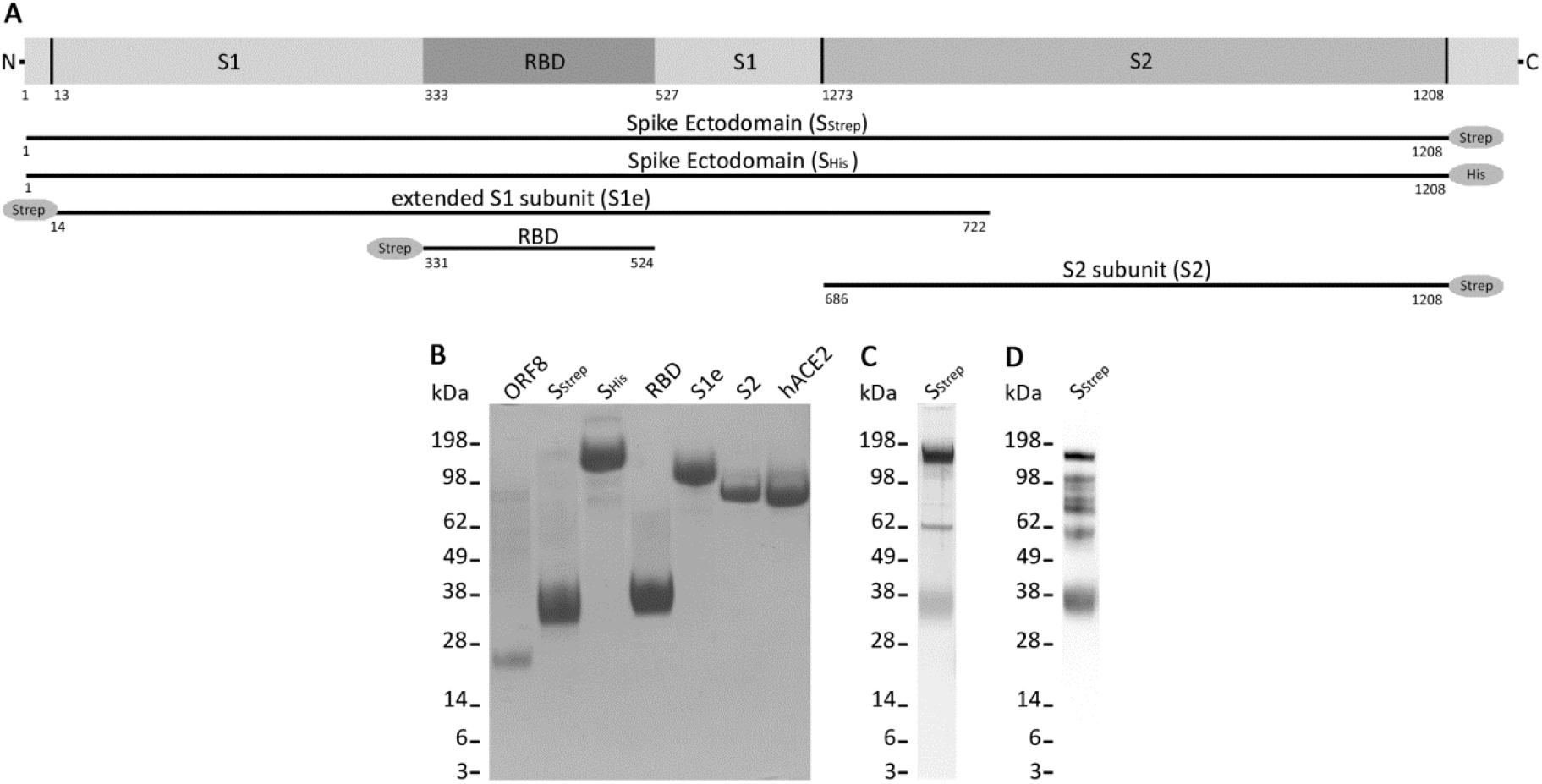
Recombinant SARS-CoV-2 proteins. (A) Schematic drawing representing recombinantly expressed SARS-CoV-2 spike glycoprotein (S_Strep_, S_His_), extended S1 subunit (S1e), receptor binding domain (RBD), and S2 subunit. (B) 4-12% Bis-Tris gel loaded with 5 μg of SARS-CoV-2 open reading frame (ORF) 8, S_Strep_, S_His_, S1e, RBD, S2, and human angiotensin converting enzyme 2 (hACE2) stained with Coomassie Brilliant Blue R 250. (C) Silver staining of 2 μg of SARS-CoV-2 S_Strep_. (D) Immunoblot of 2 μg of SARS-CoV-2 S_Strep_.

### LL-37

Synthetic LL-37 (Sequence: LLGDFFRKSKEKIGKEFKRIVQRIKDFLRNLVPRTES) was purchased from Invivogen (USA). Lyophilized peptide was reconstituted with endotoxin-free water to reach a final concentration of 1 mg/ml, which was verified by BCA assay (Thermo Fischer Scientific, USA). Purity was ≥ 95%.

### SDS-PAGE and Coomassie staining

Protein samples were reduced with dithiothreitol, mixed with LDS Sample Buffer (Thermo Fischer Scientific, USA), and heated to 90°C for 10 min. SDS-PAGE was performed in a 4-12% Bis-Tris polyacrylamide gel (Thermo Fischer Scientific, USA). Proteins were visualized using Coomassie Brilliant Blue R 250 (Merck, Germany) or using the SilverQuest silver staining kit (Invitrogen Thermo Fisher Scientific, USA). For immunoblotting, the proteins were transferred to a polyvinylidene fluoride (0.45 μm) membrane (Thermo Fischer Scientific, USA). After blocking the membrane for 1h in PBS containing 0.05% Tween and 3% BSA the membrane was incubated with Strep-Tactin^®^-HRP conjugate (IBA Lifescience, Germany) diluted 1:100,000 in blocking solution. The membrane was then treated with Amersham™ ECL™ Prime Western Blotting Detection Reagent (Thermo Fisher Scientific, USA) and signals were visualized using the ChemiDoc XRS+ System (BioRad, Germany).

### Interaction studies using surface plasmon resonance (SPR)

SPR experiments were performed as described previously ^19,20^ using a BIAcore 2000 system (BIAcore AB, Uppsala, Sweden). SARS-CoV-2 proteins were coupled to CM5 sensor chips using the amine coupling kit following the manufacturer’s instructions (Cytiva Life Sciences). SARS-CoV-2 proteins used for interaction studies were immobilized at the indicated reference units (RUs): SARS-CoV-2 ORF8 (1500 RUs), SARS-CoV-2 S_Strep_ (8000 RUs), SARS-CoV-2 S_His_ (12000 RUs), SARS-CoV-2 RBD (1500 RUs), SARS-CoV-2-S1e (6000 RUs), SARS-CoV-2 S2 (5500 RUs). Interaction studies were performed by injecting 0-1280 nM analyte in HBS-EP buffer (0.01 M HEPES, pH 7.4, 0.15 M NaCl, 3 mM EDTA, 0.005% (v/v) surfactant P20) (Cytiva Life Sciences). To verify the functionality of the ORF8 protein coupled sensor chip a Strep-Tactin-HRP conjugate (IBA Lifescience, Germany) was used as a control. Kinetic constants were calculated by nonlinear fitting (1:1 interaction model with mass transfer) to the association and dissociation curves according to the manufacturer's instructions (BIAevaluation version 3.0 software). Apparent equilibrium dissociation constants (K_D_ values) were then calculated as the ratio of the dissociation rate constant (k_d_) and the association rate constant (k_a_). Experiments were performed in quadruplicate.

### Binding competition studies using surface plasmon resonance

For competition experiments SARS-CoV-2 S_Strep_ was immobilized at 9000 RUs and 0-2560 nM LL-37 was injected followed by a subsequent 10 nM injection of hACE2 (“coinject” mode). Competition experiments with SARS-CoV-2 RBD, SARS-CoV-2 S1e, and SARS-CoV-2 S2 were performed by injecting 0-2560 nM LL-37 followed by an immediate injection of 640 nM hACE2 (“coinject” mode). Competition effects were detected as the percentage loss of signal compared to prior buffer injection.

## Results

### Recombinant expression of SARS-CoV-2 proteins

To study the interaction of LL-37 with SARS-CoV-2 we used recombinantly expressed SARS-CoV-2 S_Strep_ (139 kDa), S_His_ (136 kDa), S1e (83 kDa), RBD (26 kDa), S2 (65 kDa), and the accessory protein ORF8 (16 kDa) (Figure 1 A). Recombinant hACE2 was used as a positive control. Integrity and purity of the proteins was checked by SDS-PAGE following Coomassie staining (Figure 1 B). S_Strep_ stained with Coomassie dye showed one band with an apparent molecular mass of approximately 30 kDa and one faint band with an apparent molecular mass of 140 kDa. To verify the presence of full-length S_Strep_, we performed silver staining after SDS-PAGE, and immunoblot (Figure 1 C, D). Both showed the presence of full-length S_Strep_.

### Interaction studies using surface plasmon resonance (SPR)

To investigate the binding kinetics and interaction affinity of LL-37 with SARS-CoV-2 S protein by SPR, S_Strep_ and S_His_ were immobilized on a CM5 sensor chip (Figure 2). LL-37 showed consistent and specific binding to both spike ectodomain preparations S_Strep_ (K_D_ = 410 nM) and S_His_ (K_D_ = 420 nM) which excluded tag-specific effects. Binding of S proteins to hACE2As was measured as a positive control (Figure 2 B, D).. The binding affinities of S_Strep_ to hACE2 (K_D_ = 370 nM) and S_His_ to hACE2 (K_D_ = 370 nM) were comparable to the binding affinities seen for LL-37.

**Figure 2:**
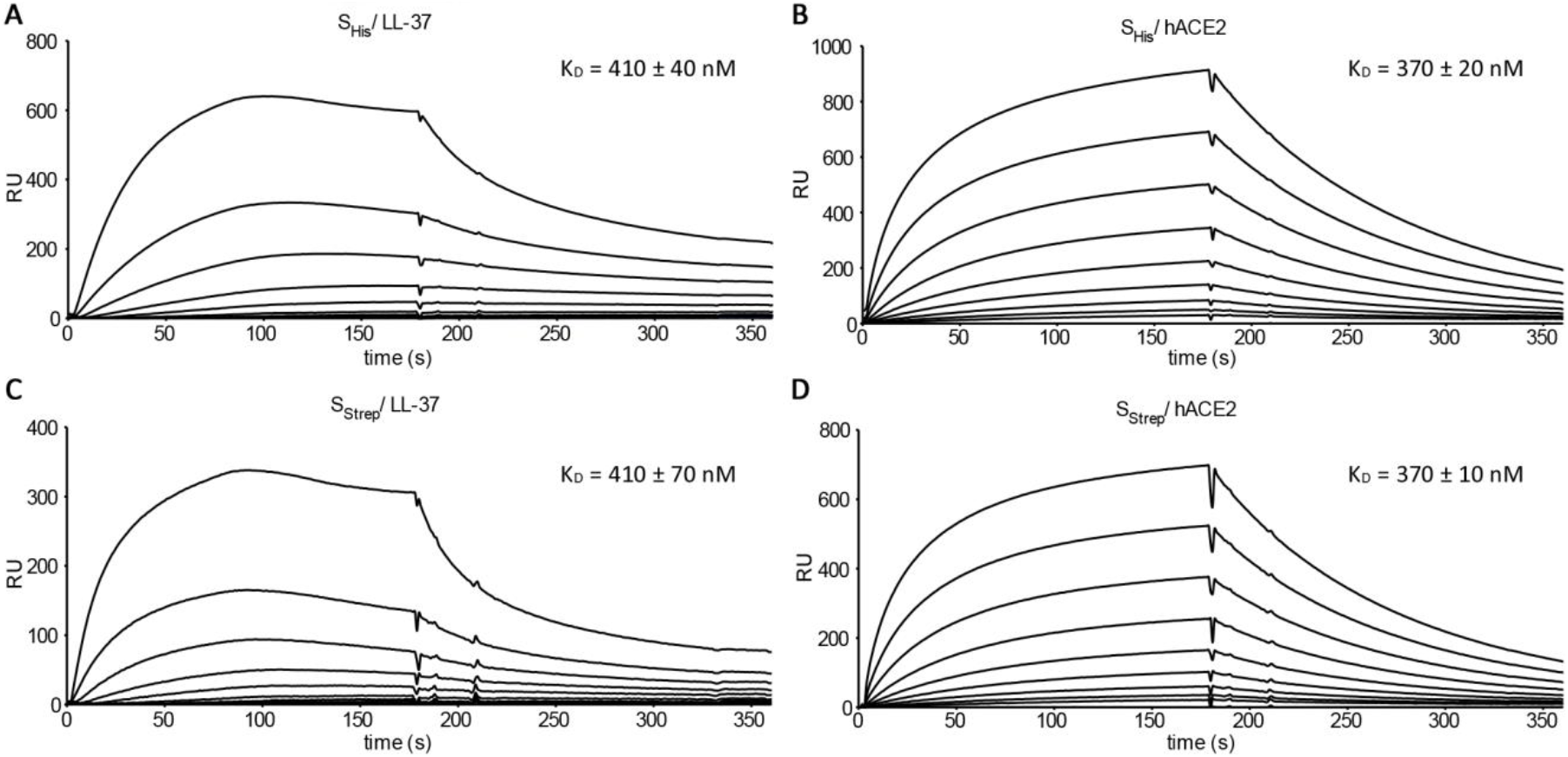
SPR sensorgrams of binding kinetics of SARS-CoV-2 S protein to LL-37 and hACE2. (A) Binding of LL-37 at various concentrations (0-1280 nM) to SARS-CoV-2 S_His_ (12000 RUs immobilized on a CM5 sensor chip) and calculated equilibrium dissociation constant (K_D_). (B) Binding of hACE2 at various concentrations (0-1280 nM) to SARS-CoV-2 S_His_ (12000 RUs immobilized on a CM5 sensor chip) and calculated equilibrium dissociation constant (K_D_). (C) Binding of LL-37 at various concentrations (0-1280 nM) to SARS-CoV-2 S_Strep_ (8000 RUs immobilized on a CM5 sensor chip) and calculated equilibrium dissociation constant (K_D_). (D) Binding of hACE2 at various concentrations (0-1280 nM) to SARS-CoV-2 S_His_ (12000 RUs immobilized on a CM5 sensor chip) and calculated equilibrium dissociation constant (K_D_).

To define the binding site of LL-37 within the S protein the recombinant proteins RBD, S1e, and S2, representing different subregions of the S protein (Figure 3), were immobilized on a CM5 sensor chip and binding of LL-37 was analyzed. LL-37 bound to RBD (K_D_ = 220 nM), S1e (K_D_ = 200 nM), and S2 (K_D_ = 230 nM) with the same binding affinities. As a control, we analyzed the binding of hACE2 to RBD (K_D_ = 940 nM), S1e (K_D_ = 610 nM) and S2. As expected, only S2, which possesses no RBD, did not show an interaction with hACE2, demonstrating the specificity of the binding of RBD and S1e to hACE2, and of RBD, S1e, and S2 to LL-37.

**Figure 3:**
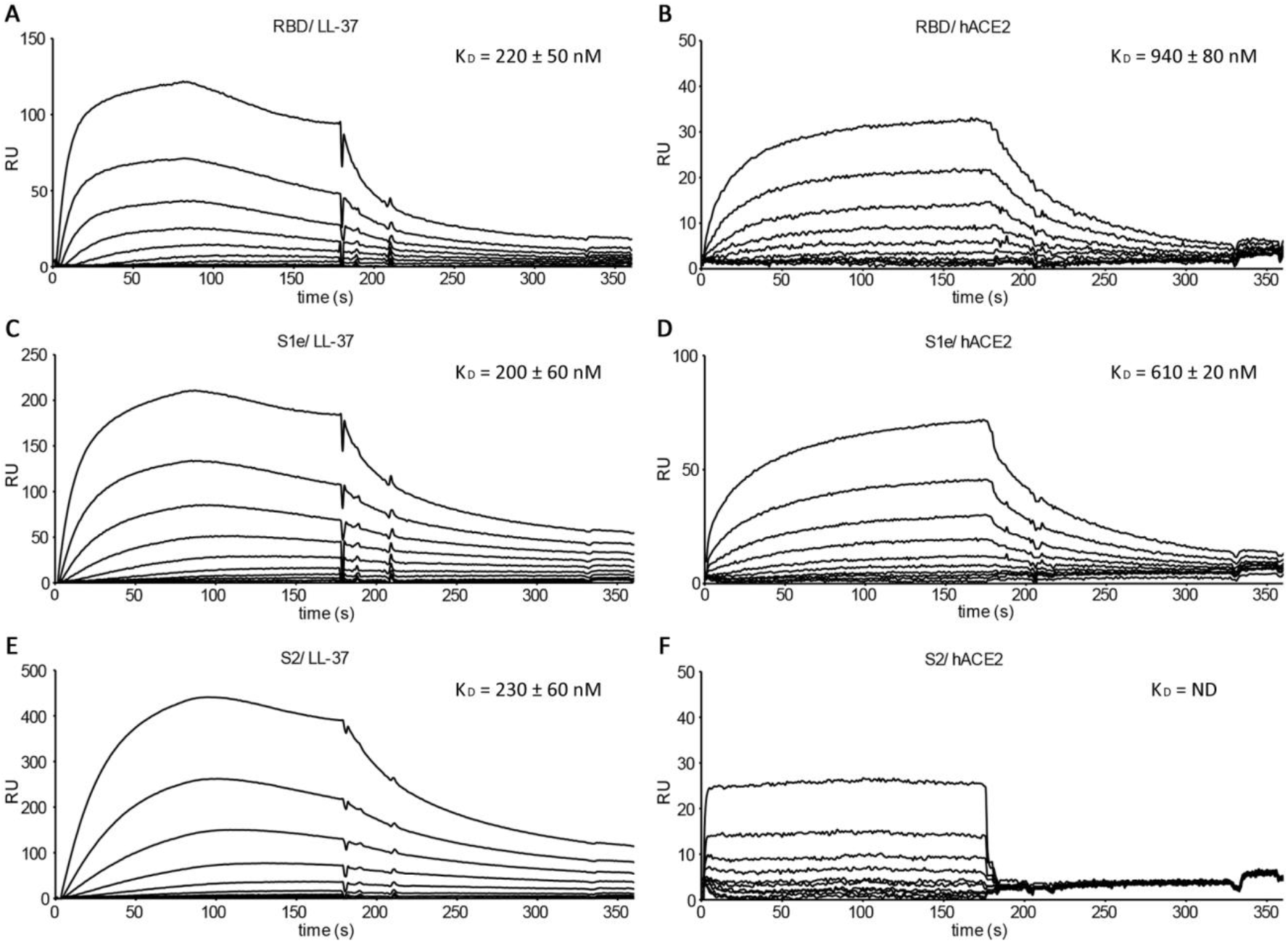
SPR sensorgrams of binding kinetics of SARS-CoV-2 RBD, S1e, and S2 to LL-37 and hACE2. (A) Binding of LL-37 at various concentrations (0-1280 nM) to SARS-CoV-2 RBD (1500 RUs immobilized on a CM5 sensor chip) and calculated equilibrium dissociation constant (K_D_). (B) Binding of hACE2 at various concentrations (0-1280 nM) to SARS-CoV-2 RBD (1500 RUs immobilized on a CM5 sensor chip) and calculated equilibrium dissociation constant (K_D_). (C) Binding of LL-37 at various concentrations (0-1280 nM) to SARS-CoV-2 S1e (6000 RUs immobilized on a CM5 sensor chip) and calculated equilibrium dissociation constant (K_D_). (D) Binding of hACE2 at various concentrations (0-1280 nM) to SARS-CoV-2 S1e (6000 RUs immobilized on a CM5 sensor chip) and calculated equilibrium dissociation constant (K_D_). (E) Binding of LL-37 at various concentrations (0-1280 nM) to SARS-CoV-2 S2 (5500 RUs immobilized on a CM5 sensor chip) and calculated equilibrium dissociation constant (K_D_). (F) Binding of hACE2 at various concentrations (0-1280 nM) to SARS-CoV-2 S2 (5500 RUs immobilized on a CM5 sensor chip) and calculated equilibrium dissociation constant (K_D_).

The accessory protein ORF8 is regarded as a virulence factor. Therefore, we also analyzed the interaction of ORF8 and LL-37 (Figure 4). Binding between LL-37 and ORF8 was detected with a K_D_ of 290 nM. A Strep-Tactin-HRP conjugate was used to verify the functionality of the ORF8 coupled sensor chip, as ORF8 carries a double Strep II tag.

**Figure 4:**
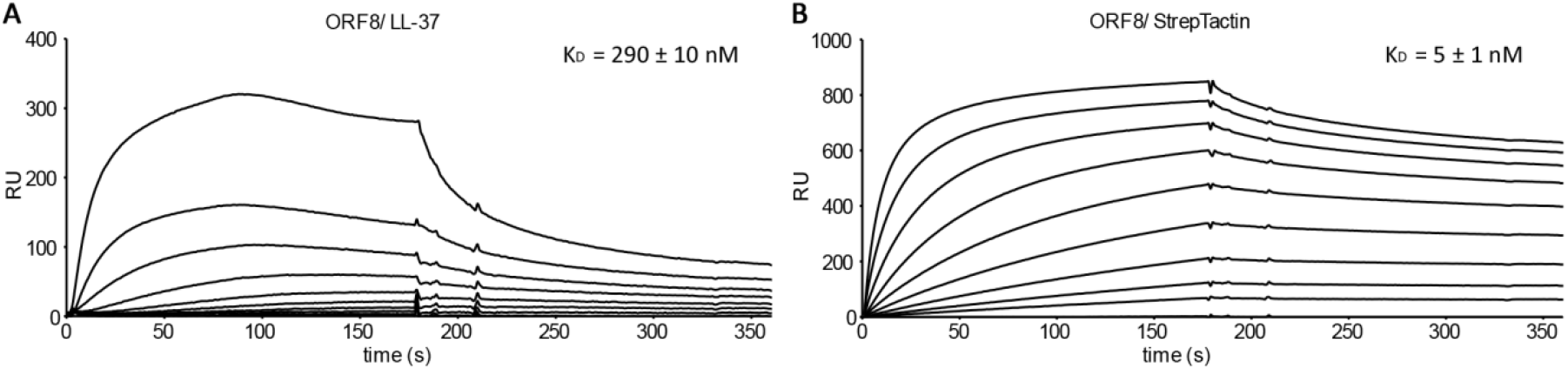
SPR sensorgrams of binding kinetics of SARS-CoV-2 ORF8 to LL-37 and Strep-Tactin-HRP conjugate. (A) Binding of LL-37 at various concentrations (0-1280 nM) to SARS-CoV-2 ORF8 (1500 RUs immobilized on a CM5 sensor chip) and calculated equilibrium dissociation constant (K_D_). (B) Binding of Strep-Tactin-HRP conjugate at various concentrations (0-1280 nM) to SARS-CoV-2 RBD (1500 RUs immobilized on a CM5 sensor chip) and calculated equilibrium dissociation constant (K_D_).

All sensorgrams of LL-37 showed a drop in the signal (RU) after approximately 100 s. This phenomenon may be caused by oligomerization which prevents further association of the monomeric analyte. It is known that LL-37 shows a concentration dependent oligomerization and forms tetrameric channels, these may in the case of bacterial infection lead to the breakdown of the transmembrane potential essential for bacterial killing ^21^.

### Binding competition studies using surface plasmon resonance

Competition studies were performed to investigate the bindingof S protein, S1e, and RBD to hACE2 by LL-37. Therefore, sensor chips were immobilized with S_Strep_, RBD, and S1e, respectively, and hACE2 was injected with a constant concentration directly after various concentrations of LL-37 (Figure 5). Calculated half maximal inhibitory concentrations (IC_50_) for the binding of S protein, S1e, and RBD to hACE2 by LL-37 were determined. LL-37 was able to reduce the binding of S_Strep_ (IC_50_ = 740 nM), S1e (IC_50_ = 170 nM), and RBD (IC_50_ = 130 nM) to hACE2. In contrast to RBD and S1e, the full-length S protein possesses additional binding sites for LL-37, which are not involved in hACE2 binding. Therefore, higher concentrations of LL-37 are needed to block the binding site of hACE2.

**Figure 5:**
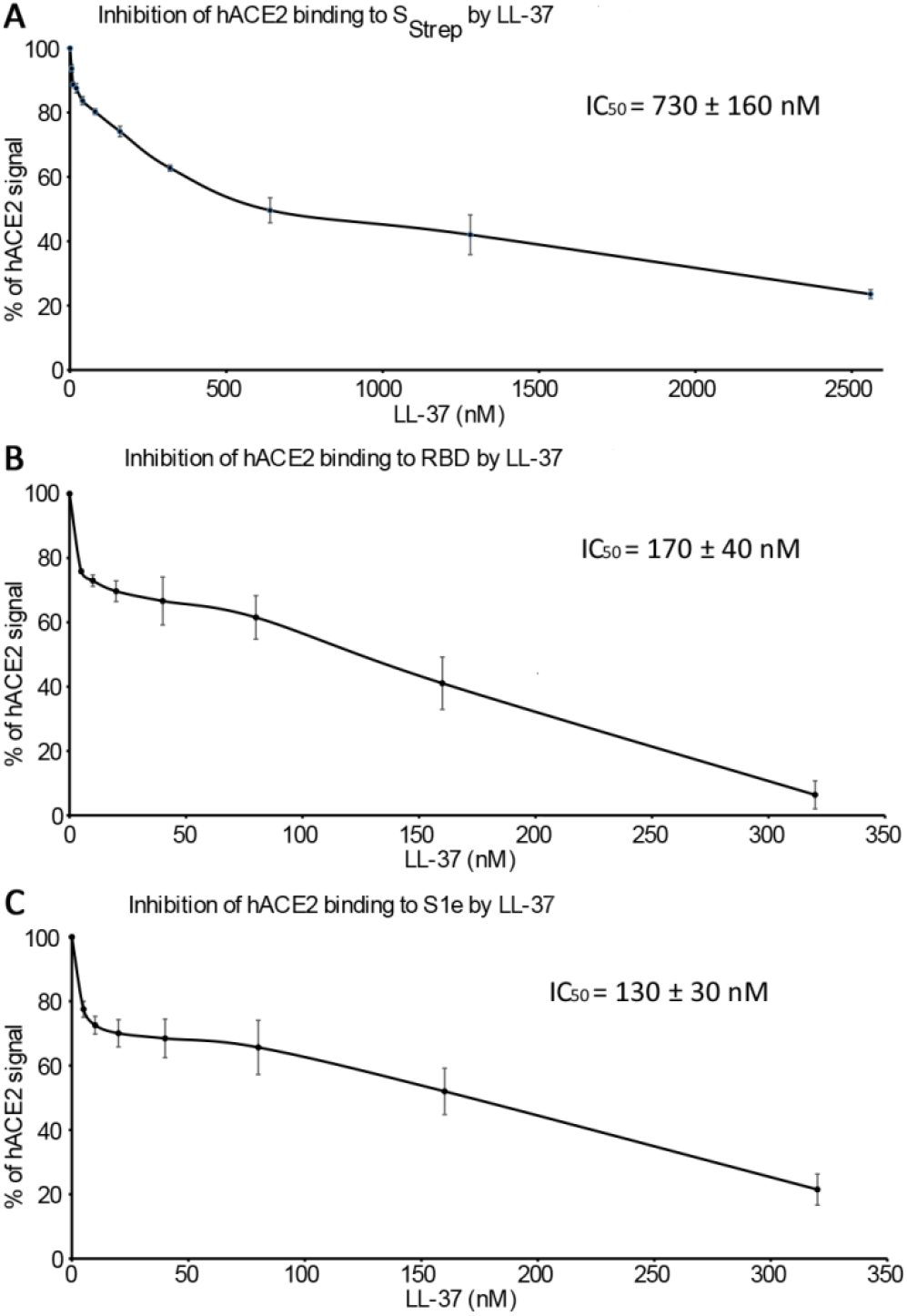
Analysis of the inhibition y LL-37 of the interactions between SARS-CoV-2 S protein, RBD, and S1e and hACE2 using competition SPR. (A) Dose response curves for IC_50_ calculation for the LL-37 inhibition of SARS-CoV-2 S_Strep_ binding to hACE2. (B) Dose response curves for IC_50_ calculation for the LL-37 inhibition of SARS-CoV-2 RBD binding to hACE2 by LL-37. (C) Dose response curves for IC_50_ calculation for the LL-37 inhibition of SARS-CoV-2 S1e binding to hACE2.

## Discussion

Several studies describe the positive effect of vitamin D on the course of COVID-19 disease, although larger independent studies are still pending. Based on the fact that gene expression of cathelicidin, the precursor of LL-37, is regulated by vitamin D, there is speculation that vitamin D has a beneficial effect on the course of COVID-19 disease through increased formation of LL-37. However, there is jet no experimental evidence for the effect of LL-37 on SARS-CoV-2 proteins. In our study, we showed for the first time a biochemical link between LL-37 and the SARS-CoV-2 S protein and provide a rationale for the use of vitamin D in infection prophylaxis, and for therapeutic administration in the treatment of COVID-19 patients. Peptide based therapeutics, especially antimicrobial peptides, are excellent drug candidates as they have little adverse effects and show anti-coronavirus activity ^22–24^. For SARS-CoV-2, our results provide evidence, that the direct use of LL-37 by inhalation or systemic application, might reduce the severity of COVID-19.

We used surface plasmon resonance analysis to show that LL-37 binds to the SARS-CoV-2 S protein and reduces the attachment of the S protein to its cellular receptor hACE2. SPR analysis demonstrated a strong interaction between LL-37 and the S protein, which most likely inhibits cellular infection. A recent study shows that at a concentration of 100 ng/ml LL-37 decreases the ability of SARS-CoV-2 to invade Vero cells by 41.5% ^25^. Our findings are in accordance with *in silico* predictions suggesting a binding of LL-37 to the RBD of SARS-CoV-2 based on a high structural similarity of LL-37 to the N-terminal helix of hACE2 ^26^. Surprisingly, LL-37 did not only bind to RBD and proteins containing the RBD, but also to S2, lacking the RBD. Therefore, we hypothesize that binding of LL-37 to the S protein does not occur through to interactions with a special protein domain, but are rather due to electrostatic or hydrophobic interactions or its amphipathicity. This assumption is further supported by the fact that the antiviral effectivity of LL-37 against influenza infection *in vitro* or *in vivo* was not altered by replacing L-amino acids with D-amino acids, indicating that its action is not dependent on specific receptor interactions ^27^. Interestingly, peptide binding to non-RBD regions can also reduce SARS-CoV-2 S protein attachment to hACE2 ^28^. Therefore, the here demonstrated binding of LL-37 to S1e and S2 may also hinder viral binding to its host receptor. In summary, we could show that LL-37 reduces attachment of the S protein to hACE2. The calculated IC_50_ values for the LL-37 inhibition of the binding of S protein and its subdomains to the hACE2 ranged from 130 nM (IC_50_ RBD) to 740 nM (IC_50_ S_Strep_). In contrast, the minimal inhibitory concentration (MIC) of LL-37 against different gram-positive and gram-negative bacteria is reported to range from 3.6 μM to 6.9 μM ^29^.

*In vitro* experiments predicted cytotoxicity of LL-37. However, in plasma up to 90% of LL-37 is bound to apolipoprotein A-I, which attenuates its cytotoxic effects *in vivo* ^30^. Thus, higher LL-37 concentrations can be reached at localized sites of infection without damaging the tissue. Within different tissues the physiological concentration of LL-37 varies. In lung and nasal secretions, high concentrations can be reached, indicating a relevant role of LL-37 in lung immune defense mechanisms ^31,32^. A study of infants hospitalized due to bronchiolitis associated low LL-37 levels with increased severity of disease ^33^. Furthermore, after cyclic stretch, typically caused by mechanical ventilation, down-regulation of the cathelicidin gene was detected in human bronchial epithelial cell lines. Treatment with vitamin D counteracted this effect on the mRNA level as well as on the protein level and protected against ventilator-induced injury ^34^. In human bronchial epithelial cells, a 10-fold induction of cathelicidin mRNA levels was achieved by vitamin D stimulation ^11^. Concerning a direct therapeutic application of antimicrobial peptides like LL-37, it is noteworthy that their pharmaceutical use has received approval for clinical trials ^25,35^. This could help to accelerate the use of LL-37 for the treatment of COVID-19.

We also tested LL-37s’ ability to bind to the negatively charged, secreted accessory protein ORF8, which is unique to SARS-CoV-2 and thus linked to efficient pathogen transmission ^36,37^. Multiple functions of ORF8 have been proposed including disruption of antigen presentation via down-regulation of MHC I expression and inhibition of interferon I signaling ^38,39^. Furthermore, a 382-nucleotide deletion in the ORF8 region of the genome led to lower concentrations of proinflammatory cytokines, chemokines, and growth factors that are strongly associated with severe COVID-19 ^2,40^. Therefore, binding of LL-37 to ORF8 may mitigate these effects leading to a less serious course of the disease.

LL-37 has not only antimicrobial and antiviral activities but also immunomodulatory properties. It helps to maintain homeostasis of the immune system by regulating chemokine, cytokine production. Immune responses are promoted through an enhanced antigen presentation by supporting the maturation of monocytes to dendritic cells, in this manner bridging innate and adaptive immunity ^41,42^. The peptide is also able to augment the release of pro-inflammatory cytokines by immunocompetent cells and human bronchial epithelial cells ^43,44^. On the other hand, it can diminish the secretion of proinflammatory factors induced by bacterial lipopolysaccharides ^45^. Moreover, influenza infected mice showed reductions of granulocyte-macrophage colony-stimulating factor (GM-CSF), RANTES, and interleukin 1ß in bronchoalveolar lavage fluid after LL-37 treatment compared to the infected control group ^27^. As severe COVID-19 patients show high levels of cytokines including IL-6, GM-CSF, RANTES, and IL-8, LL-37’s immunomodulatory properties may have protective effects and improve patient survival rates ^2,46^.

Meanwhile, vaccines against SARS-CoV-2 have been developed. With extensive vaccination the pandemic will end, but the threat of COVID-19 remains. It is for several reasons challenging to reach herd immunity. The vaccine will not be available to everyone, some individuals have contraindications or are not willing to be vaccinated and others will not develop effective immunity after vaccination. For those patients, prophylactic and therapeutic agents directed against COVID-19 are urgently needed. The combined antiviral, antimicrobial, and immunomodulatory properties are advantages of LL-37 compared to antibodies from reconvalescent donors and should be considered as COVID-19 patients are prone to develop bacterial superinfections. Furthermore, the number of plasma donors limits the quantity of reconvalescent plasma, and the transfer of human material carries the risk of infection. In contrast, LL-37 can be produced synthetically with high purity and in unlimited quantities. Also, the use of antibodies from reconvalescent donors did not result in clinical benefits or in reduction in all-cause mortality ^47,48^. Therefore, the use of vitamin D as an infection prophylaxis and the therapeutic administration of vitamin D for the treatment of COVID-19 patients, and the direct use of LL-37 by inhalation or systemic application could be good alternatives.

In conclusion, we have revealed a biochemical link between vitamin D, LL-37, and COVID-19 severity. SPR analysis demonstrated that LL-37 binds to SARS-CoV-2 S protein and inhibits binding to its receptor hACE2, and most likely viral entry into the cell. This study supports the prophylactic use of vitamin D to induce LL-37 to protect from SARS-CoV-2 infection, and the therapeutic administration of vitamin D for the treatment of COVID-19 patients. Further, our results provide evidence that the direct use of LL-37 by inhalation and systemic application might reduce the severity of COVID-19.

## Acknowledgements

Funding for this study was provided by the Deutsche Forschungsgemeinschaft (DFG) via FOR2722/ C2 to G.S. and FOR2722/ B2 to M.K..

## Author Contributions

Pending

## Literature

1 Perlman, S. & Netland, J. Coronaviruses post-SARS: update on replication and pathogenesis. Nature Reviews Microbiology 7, 439–450, doi:10.1038/nrmicro2147 (2009).

2 Huang, C. et al. Clinical features of patients infected with 2019 novel coronavirus in Wuhan, China. The Lancet 395, 497–506, doi:https://doi.org/10.1016/S0140-6736(20)30183-5 (2020).

3 Ksiazek, T. G. et al. A Novel Coronavirus Associated with Severe Acute Respiratory Syndrome. New England Journal of Medicine 348, 1953–1966, doi:10.1056/NEJMoa030781 (2003).

4 Zaki, A. M., van Boheemen, S., Bestebroer, T. M., Osterhaus, A. D. M. E. & Fouchier, R. A. M. Isolation of a Novel Coronavirus from a Man with Pneumonia in Saudi Arabia. New England Journal of Medicine 367, 1814–1820, doi:10.1056/NEJMoa1211721 (2012).

5 Li, H. et al. Updated Approaches against SARS-CoV-2. Antimicrobial agents and chemotherapy 64, doi:10.1128/aac.00483-20 (2020).

6 Rhodes, J. M., Subramanian, S., Laird, E., Griffin, G. & Kenny, R. A. Perspective: Vitamin D deficiency and COVID-19 severity - plausibly linked by latitude, ethnicity, impacts on cytokines, ACE2 and thrombosis. J Intern Med, doi:10.1111/joim.13149 (2020).

7 Entrenas Castillo, M. et al. “Effect of calcifediol treatment and best available therapy versus best available therapy on intensive care unit admission and mortality among patients hospitalized for COVID-19: A pilot randomized clinical study”. The Journal of Steroid Biochemistry and Molecular Biology 203, 105751, doi:https://doi.org/10.1016/j.jsbmb.2020.105751 (2020).

8 Bleizgys, A. Vitamin D and COVID-19: It is time to act. International Journal of Clinical Practice, e13748, doi:https://doi.org/10.1111/ijcp.13748 (2020).

9 Miller, J. & Gallo, R. L. Vitamin D and innate immunity. Dermatologic Therapy 23, 13–22, doi:10.1111/j.1529-8019.2009.01287.x (2010).

10 Wang, T. T. et al. Cutting edge: 1,25-dihydroxyvitamin D3 is a direct inducer of antimicrobial peptide gene expression. J Immunol 173, 2909–2912, doi:10.4049/jimmunol.173.5.2909 (2004).

11 Yim, S., Dhawan, P., Ragunath, C., Christakos, S. & Diamond, G. Induction of cathelicidin in normal and CF bronchial epithelial cells by 1,25-dihydroxyvitamin D(3). Journal of cystic fibrosis : official journal of the European Cystic Fibrosis Society 6, 403–410, doi:10.1016/j.jcf.2007.03.003 (2007).

12 Gombart, A. F., Borregaard, N. & Koeffler, H. P. Human cathelicidin antimicrobial peptide (CAMP) gene is a direct target of the vitamin D receptor and is strongly up-regulated in myeloid cells by 1,25-dihydroxyvitamin D3. FASEB journal : official publication of the Federation of American Societies for Experimental Biology 19, 1067–1077, doi:10.1096/fj.04-3284com (2005).

13 Liu, P. T. et al. Convergence of IL-1beta and VDR activation pathways in human TLR2/1-induced antimicrobial responses. PloS one 4, e5810, doi:10.1371/journal.pone.0005810 (2009).

14 Pahar, B., Madonna, S., Das, A., Albanesi, C. & Girolomoni, G. Immunomodulatory Role of the Antimicrobial LL-37 Peptide in Autoimmune Diseases and Viral Infections. Vaccines 8, doi:10.3390/vaccines8030517 (2020).

15 Beard, J. A., Bearden, A. & Striker, R. Vitamin D and the anti-viral state. Journal of Clinical Virology 50, 194–200, doi:10.1016/j.jcv.2010.12.006 (2011).

16 Dömling, A. & Gao, L. Chemistry and Biology of SARS-CoV-2. Chem 6, 1283–1295, doi:https://doi.org/10.1016/j.chempr.2020.04.023 (2020).

17 Tan, Y., Schneider, T., Leong, M., Aravind, L. & Zhang, D. Novel Immunoglobulin Domain Proteins Provide Insights into Evolution and Pathogenesis of SARS-CoV-2-Related Viruses. mBio 11, e00760–00720, doi:10.1128/mBio.00760-20 (2020).

18 Kowarz, E., Löscher, D. & Marschalek, R. Optimized Sleeping Beauty transposons rapidly generate stable transgenic cell lines. Biotechnology journal 10, 647–653, doi:10.1002/biot.201400821 (2015).

19 Sengle, G. et al. Targeting of bone morphogenetic protein growth factor complexes to fibrillin. The Journal of biological chemistry 283, 13874–13888, doi:10.1074/jbc.M707820200 (2008).

20 Sengle, G., Ono, R. N., Lyons, K. M., Bächinger, H. P. & Sakai, L. Y. A new model for growth factor activation: type II receptors compete with the prodomain for BMP-7. J Mol Biol 381, 1025–1039, doi:10.1016/j.jmb.2008.06.074 (2008).

21 Sancho-Vaello, E. et al. The structure of the antimicrobial human cathelicidin LL-37 shows oligomerization and channel formation in the presence of membrane mimics. Scientific Reports 10, 17356, doi:10.1038/s41598-020-74401-5 (2020).

22 Guo, Y. et al. Identification of a New Region of SARS-CoV S Protein Critical for Viral Entry. J Mol Biol 394, 600–605, doi:https://doi.org/10.1016/j.jmb.2009.10.032 (2009).

23 Sagar, S. et al. Bromelain Inhibits SARS-CoV-2 Infection in VeroE6 Cells. bioRxiv : the preprint server for biology, 2020.2009.2016.297366, doi:10.1101/2020.09.16.297366 (2020).

24 Zhang, G. et al. Investigation of ACE2 N-terminal fragments binding to SARS-CoV-2 Spike RBD. bioRxiv, 2020.2003.2019.999318, doi:10.1101/2020.03.19.999318 (2020).

25 Zhang, H. et al. Preliminary evaluation of the safety and efficacy of oral human antimicrobial peptide LL-37 in the treatment of patients of COVID-19, a small-scale, single-arm, exploratory safety study. medRxiv, 2020.2005.2011.20064584, doi:10.1101/2020.05.11.20064584 (2020).

26 Lokhande, K. B., Banerjee, T., Venkateswara, S., K & Deshpande, M. An in Silico Scientific Basis for LL-37 as a Therapeutic and Vitamin D as Preventive for Covid-19. chemRxiv, doi:10.26434/chemrxiv.12928202 (2020).

27 Barlow, P. G. et al. Antiviral activity and increased host defense against influenza infection elicited by the human cathelicidin LL-37. PloS one 6, e25333, doi:10.1371/journal.pone.0025333 (2011).

28 Qiao, B. & Olvera de la Cruz, M. Enhanced Binding of SARS-CoV-2 Spike Protein to Receptor by Distal Polybasic Cleavage Sites. ACS Nano 14, 10616–10623, doi:10.1021/acsnano.0c04798 (2020).

29 Bals, R., Wang, X., Zasloff, M. & Wilson, J. M. The peptide antibiotic LL-37/hCAP-18 is expressed in epithelia of the human lung where it has broad antimicrobial activity at the airway surface. Proc Natl Acad Sci U S A 95, 9541–9546, doi:10.1073/pnas.95.16.9541 (1998).

30 Svensson, D., Lagerstedt, J. O., Nilsson, B. O. & Del Giudice, R. Apolipoprotein A-I attenuates LL-37-induced endothelial cell cytotoxicity. Biochemical and biophysical research communications 493, 71–76, doi:10.1016/j.bbrc.2017.09.072 (2017).

31 Kim, S. T. et al. Antimicrobial peptide LL-37 is upregulated in chronic nasal inflammatory disease. Acta oto-laryngologica 123, 81–85, doi:10.1080/0036554021000028089 (2003).

32 Schaller-Bals, S., Schulze, A. & Bals, R. Increased levels of antimicrobial peptides in tracheal aspirates of newborn infants during infection. American journal of respiratory and critical care medicine 165, 992–995, doi:10.1164/ajrccm.165.7.200110-020 (2002).

33 Mansbach, J. M. et al. Serum LL-37 Levels Associated With Severity of Bronchiolitis and Viral Etiology. Clinical infectious diseases : an official publication of the Infectious Diseases Society of America 65, 967–975, doi:10.1093/cid/cix483 (2017).

34 Karadottir, H., Kulkarni, N. N., Gudjonsson, T., Karason, S. & Gudmundsson, G. H. Cyclic mechanical stretch down-regulates cathelicidin antimicrobial peptide expression and activates a pro-inflammatory response in human bronchial epithelial cells. PeerJ 3, e1483, doi:10.7717/peerj.1483 (2015).

35 Koo, H. B. & Seo, J. Antimicrobial peptides under clinical investigation. Peptide Science 111, e24122, doi:https://doi.org/10.1002/pep2.24122 (2019).

36 Wang, X. et al. Accurate Diagnosis of COVID-19 by a Novel Immunogenic Secreted SARS-CoV-2 orf8 Protein. mBio 11, e02431–02420, doi:10.1128/mBio.02431-20 (2020).

37 Hassan, S. S. et al. A unique view of SARS-CoV-2 through the lens of ORF8 protein. bioRxiv, 2020.2008.2025.267328, doi:10.1101/2020.08.25.267328 (2020).

38 Zhang, Y. et al. The ORF8 Protein of SARS-CoV-2 Mediates Immune Evasion through Potently Downregulating MHC-I. bioRxiv, 2020.2005.2024.111823, doi:10.1101/2020.05.24.111823 (2020).

39 Li, J. Y. et al. The ORF6, ORF8 and nucleocapsid proteins of SARS-CoV-2 inhibit type I interferon signaling pathway. Virus research 286, 198074, doi:10.1016/j.virusres.2020.198074 (2020).

40 Young, B. E. et al. Effects of a major deletion in the SARS-CoV-2 genome on the severity of infection and the inflammatory response: an observational cohort study. Lancet (London, England) 396, 603–611, doi:10.1016/s0140-6736(20)31757-8 (2020).

41 Bandholtz, L. et al. Antimicrobial Peptide LL-37 Internalized by Immature Human Dendritic Cells Alters their Phenotype. Scandinavian Journal of Immunology 63, 410–419, doi:https://doi.org/10.1111/j.1365-3083.2006.001752.x (2006).

42 Davidson, D. J. et al. The cationic antimicrobial peptide LL-37 modulates dendritic cell differentiation and dendritic cell-induced T cell polarization. J Immunol 172, 1146–1156, doi:10.4049/jimmunol.172.2.1146 (2004).

43 Pistolic, J. et al. Host defence peptide LL-37 induces IL-6 expression in human bronchial epithelial cells by activation of the NF-kappaB signaling pathway. J Innate Immun 1, 254–267, doi:10.1159/000171533 (2009).

44 Yang, B. et al. Significance of LL-37 on Immunomodulation and Disease Outcome. Biomed Res Int 2020, 8349712–8349712, doi:10.1155/2020/8349712 (2020).

45 Mookherjee, N. et al. Modulation of the TLR-Mediated Inflammatory Response by the Endogenous Human Host Defense Peptide LL-37. The Journal of Immunology 176, 2455–2464, doi:10.4049/jimmunol.176.4.2455 (2006).

46 Tang, Y. et al. Cytokine Storm in COVID-19: The Current Evidence and Treatment Strategies. Front Immunol 11, 1708–1708, doi:10.3389/fimmu.2020.01708 (2020).

47 Agarwal, A. et al. Convalescent plasma in the management of moderate covid-19 in adults in India: open label phase II multicentre randomised controlled trial (PLACID Trial). BMJ 371, doi:10.1136/bmj.m3939 (2020).

48 Pathak, E. B. Convalescent plasma is ineffective for covid-19. BMJ 371, m4072, doi:10.1136/bmj.m4072 (2020).

